# Cable Energy Function of Cortical Axons: Equivalent Formulas

**DOI:** 10.1101/341701

**Authors:** Zhe Jiao, Yuguo Yu

**Affiliations:** State Key Laboratory of Medical Neurobiology, School of Life Science and Human Phenome Institute, Institutes of Brain Science, Institute of Science and Technology for Brain-Inspired Intelligence, Fudan University, Shanghai, China 200433; Department of Applied Mathematics, Northwestern Polytechnical University, Xi’an 710129, People’s Republic of China

**Keywords:** energy consumption, neuron, cortical axons, Hodgkin-Huxley equation, Computational model

## Abstract

Cortical neurons generally have rich morphologies in dendrite arbor and axonal branches, which make it difficulty in estimate energy consumption during action potential (AP) propagation in neuronal communication. It is an unsolved issue in driving general analytical equations to estimate energy cost for those axons and dendrites with different terminations. Most previous energy calculations of AP-related metabolic cost are still based on the Na ^+^ -counting method. Here, we apply principles of physics and mathematical analysis to construct several forms of cable energy function of AP conduction along axons with different boundary conditions. These derived energy equations extend Hodgkin-Huxley theory and prove to be highly more accurate in estimation the energy consumption during AP propagation along cortical axons and dendrites with any kind of ion channels than that using the Na ^+^ -counting method.

**Summary:** Accurate energy estimation of action potential conduction along axons with different complex terminal conditions is an unsolved issue. We have applied principles of physics and mathematical analysis to derive several forms of cable energy function of action potential conduction along cortical axons with different boundary conditions, and we have proved that these functions are equivalent. The energy calculations of action potential metabolic cost by using our cable energy function is proved to be highly accurate than that based on the Na ^+^ -counting method. This mathematical framework allows us to estimate the energy used by AP propagation along cortical axons and dendrites with any kind of ion channels more accurately than that using the Na ^+^ -counting method. Accurate calculation of energy consumption of AP conduction may be crucial in the estimation of energy expenditure, from subcellular to whole-brain level. In addition, the analytical formula of energy calculation is valuable in investigating the key factors that influence energy consumption and reveal trade-offs between energetic constraints and neural coding efficiency for individual neurons with rich morphology structures.

## Introduction

Accurate estimation of the metabolic cost of action potential generation and propagation is important for the calculation of energy budgets for individual neurons as well as for the whole brain (Hofman, 1983;Attwell and Laughlin, 2001;Lennie, 2003;Henry, 2005;Alle et al., 2009;Carter and Bean, 2009;Herman et al., 2009;Hasenstaub et al., 2010;Sengupta et al., 2010;Belanger et al., 2011;Harris and Attwell, 2012;Howarth et al., 2012;Yu et al., 2012;Hyder et al., 2013b;Moujahid et al., 2014;Sanganahalli et al., 2016). These estimates reveal computational rules such as optimal trade-offs between metabolic constraints and neural coding (Laughlin, 2001;Laughlin and Sejnowski, 2003;Lennie, 2003;Moujahid et al., 2011;Sengupta et al., 2013) and improve the interpretation of functional magnetic resonance imaging data(Logothetis, 2008;Hyder et al., 2013a;Hyder et al., 2013b;Shulman et al., 2014;Magistretti and Allaman, 2015;Yu et al., 2017).

In previous studies, estimating the metabolic cost of APs was usually done by recording sodium currents underlying action potentials and calculating the total amount of sodium ions for each AP (Attwell and Laughlin, 2001;Laughlin, 2001;Lennie, 2003;Carter and Bean, 2009;Sengupta et al., 2010;Hallermann et al., 2012;Harris and Attwell, 2012). The total Na ^+^ provides an estimate of the number of pump cycles or ATP molecules that the active Na^+^/K^+^ pump needs to re-establish the resting state of the neuron. However, the Na ^+^ -counting method is far from accurate because it does not include all of the factors consuming ATP, thus seriously underestimating the metabolic costs of AP generation (Ju et al., 2016).

To address these shortcoming, energy estimation based on the electrochemical energy function was first performed using a single-compartment Hodgkin-Huxley neuron model(Moujahid et al., 2011;Moujahid and d’Anjou, 2012;Moujahid et al., 2014). Ju et al. developed an energy function for cortical axons (Ju et al., 2016). The analytical approach of the axonal energy function provided an inhomogeneous distribution of metabolic cost along an axon with either uniformly or nonuniformly distributed ion channels. Their results showed that the Na+-counting method severely underestimates energy cost in the cable model by 20–70%. However, this previous analytical work did not satisfactorily consider boundary conditions, as real neuronal axons and dendrites generally have branch point or sealed ends. In this paper, we comprehensively investigate how to calculate the electrochemical energy for a cable equation modeling the cortical axon and derive several equivalent energy functions associated with AP propagation under some assumptions of axon terminations. Then, we compared these results with previous works.

## Results

### The Cable Energy Function for a Cortical Axon

To precisely estimate the total amount of energy associated with AP generation and propagation along neuronal axons, we refer to the well-developed cable theory (Rall, 1989) that has been shown to be consistent with experimental observations(Shu et al., 2006;McCormick et al., 2007;Shu et al., 2007;Yu et al., 2008). Multiplying both sides of the cable equation ([M6]) (see Methods section) by the membrane potential *V*(*x*,*t*) and integrating from 0 to *t* for the axon from 0 to end *l*, the energy consumption of the whole axonal electronic circuit is derived as the following
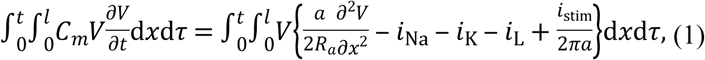

where *C_m_* is the membrane capacitance, *a* is the axonal radius, *R_a_* is the axial intracellular resistivity in Ohms centimeters, and *i*_stim_(*x,t*) is the AP-stimulating current in micro-amp per centimeter.

Based on the cable equation that describes how ion currents flow along the cable as well as analysis of the electrical energy in the equivalent circuit, we derive the first energy function of the total electrochemical energy cost of AP propagation in the given cortical axon as
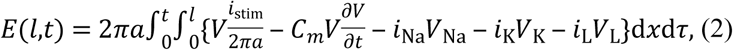

the first term of which represents the electrical energy given to the axon at some site, and the other four terms of which represent the total electrical energy accumulated in the equivalent circuit at a time interval (0,*t*). From Eq. [1] and Eq. [2], we obtain
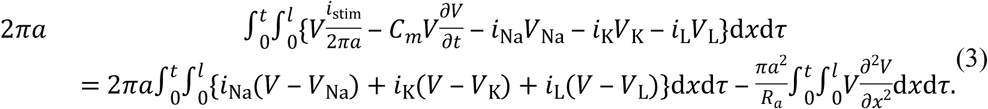

Notably, the right-hand side of Eq. [3]
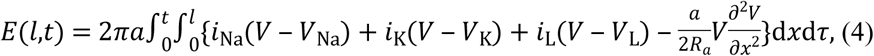

is the same energy function as that derived in our previous work (Ju et al., 2016). Here, we do not add any assumption on the axon terminations; therefore, the energy functions Eq. [2] and Eq. [4] are both applicable to any type of cortical cable model with different terminations (see Methods). In previous work (Ju et al., 2016), the authors derived Eq. [4] based on the method used in previous references(Moujahid et al., 2011), the viewpoints of which are totally different from ours here.

### The Cable Energy Function for a Cortical Axon with Different Terminations

Next, we applied the above energy function to study the energy function for a cortical cable with killed or sealed terminations. Integrating by parts, we know that the last term of Eq. [3] is
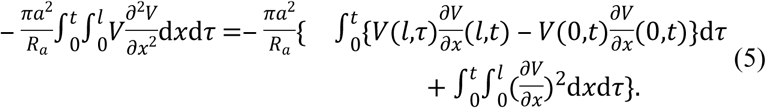

For the ends of the cortical axon that are sealed or killed, by the given boundary conditions, we obtain
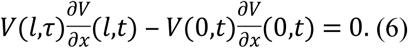

From ([4])-([6]), we have another equivalent energy function
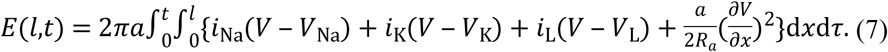

From the mathematical calculation, we know that energy functions Eq. [2], Eq. [4] and Eq. [7] are analytically equivalent in this case of boundary conditions, which is also proven by the numerical results (see Table [T2] and Fig. [fig: 1]D). Furthermore, when we compared the last term of Eq. [4] with Eq. [7], we have
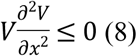

in the case of the cortical axon with terminated or sealed ends. For a given site 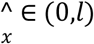, the inequality Eq. [8] implies that the solution 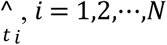 (*N* is obviously an even number), to 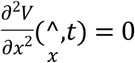 are *inflection points*, and 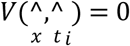. Immediately, we know that the firing rate at 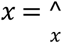 is 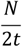. Moreover, from the property of the inflection points, we know that the curve of action potential 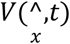between *t*_2*k* - 1_ and *t*_2*k*_, 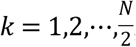 is *convex*, and the other part of the curve is *concave*.

**Figure 1.**
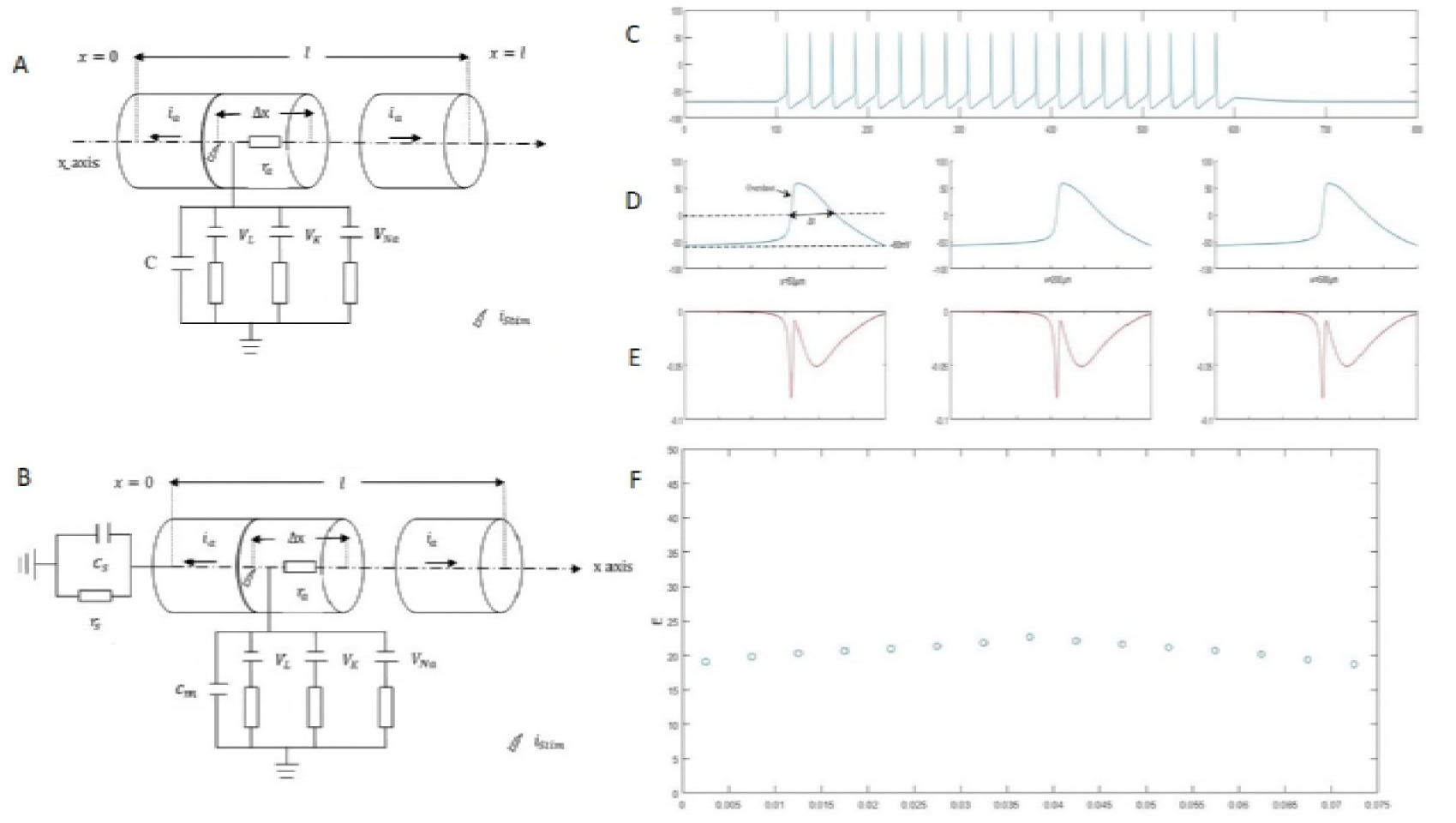
(A) Cable model of a cortical axon with its terminations killed or sealed. (B) Cable model of a cortical axon with its terminations connecting with a soma. All the parameters of the cortical cable models are given in Table [T1]. (C) APs (*N* = 40, 40*Hz*) are initiated by a stimulating current *i*(*t*) at *x* = 400 *μm*. (D) Traces of APs in the compartments of a uniform axon as in the model A (*x* = 50 *μm*, *x* = 200 *μm* and *x* = 500 *μm*). The overshoot part of the APs (*V*(*x*,*t*) > 0) is convex, and the other part is concave. can replace the full width at half maximum to study the neural coding. (E) Traces of the total energy consuming rate (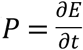) in the compartments at *x* = 50 *μm*, *x*=200 *μm* and *x* = 500 *μm*. Since the cortical energy functions of [2], [4] and [7] are equivalent, the values of the energy consuming rates are equal to one another. (F) Distribution of energy cost per unit membrane area at different compartments of the axon. The distribution is calculated by using the cable energy function [7]. Note that the same distributions can be obtained by using the cable energy function [2] and [7].

If the end *x* = 0 of the axon connects to the soma (lumped-soma termination, see Methods) and the other end is killed or sealed, that is, *V*(*l*,*τ*) = 0 or 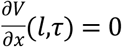 and 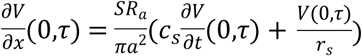 then we have
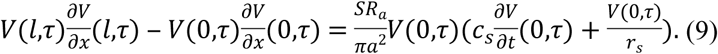

Assuming the cortical axon returns to the initial state at time, we have *V*(0,*t*) = *V*(0,0) =–70 mV. From Eq. [8] and (Methods Equation [M7]) we have
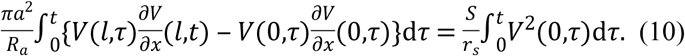

By Eqs. [4], [5] and [9], we have the following energy function
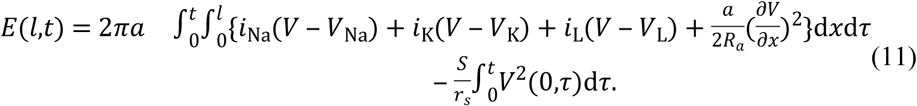

As above, we see that the energy functions Eqs. [2], [4] and [10] are analytically equivalent if one axon end (x = l) is sealed or killed and the other axon end (x = 0) connects to the spherical soma. Therefore, the energy cost of AP propagation is obviously less than the cost in other cases.

## Discussion

Based on mathematical analysis and physical intuition, we systematically establish the energy-function method for the cable model of cortical axons. In this paper, we focus on the analytical approach, which is more mathematically accurate. By the analytical equivalence of Eq. [2] and Eq. [4], we can obtain identical numerical results as previous work (Ju et al., 2016) showing the following: 1. AP propagation requires more energy than the amount estimated by the point model, and 2. the energy consumption rate of the entire branched axon scales at 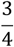 power of axonal volume. See Results for details.

The cable energy functions derived in this paper provide an accurate and applicable way to estimate the energy consumption of electric signals conducted in any type of cortical axon with different terminations. This mathematical framework allows us to estimate the energy used by AP propagation along cortical axons and dendrites with any kind of ion channels more accurately than that using the Na ^+^ -counting method. Accurate calculation of energy consumption of AP conduction may be crucial in the estimation of energy expenditure, from subcellular to whole-brain level. In addition, this analytical method of energy calculation can be used to investigate factors that influence energy consumption and reveal trade-offs between energetic constraints and neural coding efficiency for individual neurons with rich morphology structures.

## Methods

### Cable Model of the Cortical Axon

To describe AP propagation along an axon, the cable equation that describes the flow of ion currents along the axon needs to be derived (Gabbiani and Cox, 2010). Figure [fig: 1] A and B give the equivalent circuit of the cable model. During AP propagation, the membrane potential, *V*(*x*,*t*), changes along the *x*-axis, and a longitudinal current, *i*(*x*,*t*), passing through causes a voltage drop across the longitudinal resistor, *R_L_*. According to Ohm’s law, we have
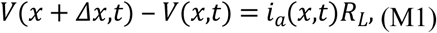

where *R_L_* = *r_a_*Δ*x*= with 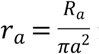 is in units of Ohms per centimeter, and *R_a_* is the axial intracellular resistivity in Ohms centimeters. For the transversal current, Kirchhoff’ s law regarding the conservation of current at each node leads to
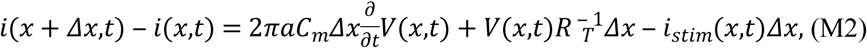

where *C_m_* is in Farads per square centimeter, *R_m_* is the resistivity of the membrane material, 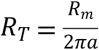 is in Ohms centimeters, *R_m_* is in Ohms square centimeters, and *i*_stim_(*x*,*t*) is the AP-stimulating current in micro-amp per centimeter. Thus, taking *Δx*→0 in [M1] and [M2], we have

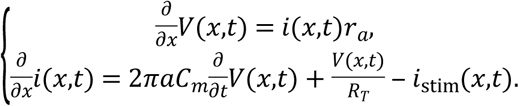

Then, we have the linear cable equation
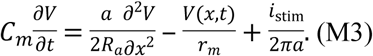

In the context of a realistic modeling of cortical neurons, we can add ion channels to the circuit diagram. Then, in the equation [M3], the term 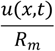 should be replaced by

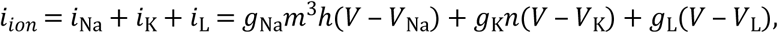

where *g*_Na_, *g*_K_, and *g*_L_, and are the maximal sodium, maximal potassium, and leak conductance per unit membrane area, respectively; and *V*_Na_, *V*_K_, and *V*_L_ are the reversal potentials of the sodium, potassium, and leak channels, respectively. The gate variables,*m*, *h*, and *n*, are dimensionless activation and inactivation variables, which describe the activation and inactivation process of the sodium and potassium channels and are governed by the following differential equations
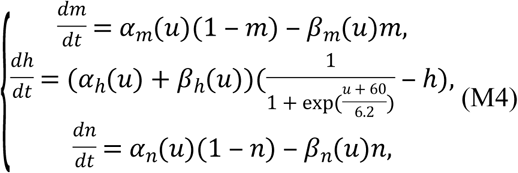

with the functions
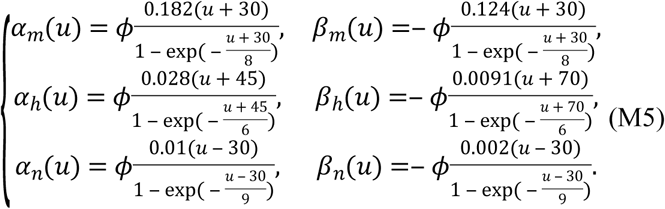

Here, 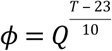 regulates the temperature dependence. Thus, we obtain the cortical cable equation
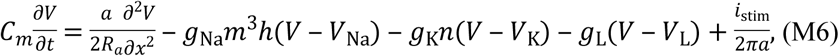

where *m*, *h*, and *n* are governed by [M4] and [M5]. We assume that the neuron starts form rest.

That is, the initial data *V*(*x*,0) =– 70 mV. We input the following form of stimulus to the cable.

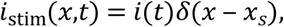

where *δ*(*x*) is the Dirac’s function, *x*_s_ ∊ (0,*l*) and

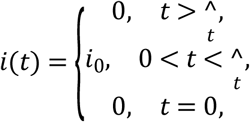

where *i*_0_ is larger than the threshold. Therefore, we input the initial threshold at the interval 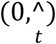 and the site *x* = *x_s_* of the cable.

We consider each of the following boundary conditions. *Sealed end*: the sealed end, which is sometimes referred to as an open-circuit termination, means that there is no longitudinal current at the end. If the longitudinal current is zero, the boundary condition is 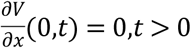 or if applied at *x* = *l*, 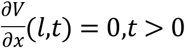. *Killed end*: a killed end, which is sometimes called a short-circuit termination, refers to the sudden ending of the nerve-cylinder membrane without a terminal covering membrane, which causes the intracellular fluid to end abruptly and abut the extracellular fluid. Thus, the appropriate boundary condition is *V*(0,*t*) = 0, *t* > 0, or if applied at *x* = *l*, *V*(*l*,*t*) = 0,*t* > 0. *Lumped-soma termination*: a lumped soma refers to the treatment of the soma as an equipotential surface, where *S* is the somatic membrane area, and it is regarded as a single resistance, *r_s_*, and capacitance, *C_s_*, attached to a nerve cylinder as in the case of a natural termination. Then, the boundary condition for a lumped soma at *x* = 0 is (see Fig. [fig:1]B)

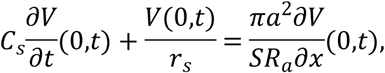

and if the end at *x* = *l* is sealed, 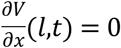

All the parameters (see Table [T1]) in the paper are within the physiological regime(Koch and HC/Biologie, 1999).

**Table 1.** Parameters of the Cable equations.

|  |  |
| --- | --- |
| $C_m$ | $0.75 \mu\text{F} \cdot \text{cm}^{-2}$ |
| $C_s$ | $0.75 \mu\text{F} \cdot \text{cm}^{-2}$ |
| $a$ | $0.75 \mu\text{m}$ |
| $R_a$ | $0.15 \text{k}\Omega \cdot \text{cm}$ |
| $r_s$ | $0.1 \text{k}\Omega \cdot \text{cm}$ |
| $T$ | $37^\circ \text{C}$ |
| $Q$ | $2.3$ |
| $S$ | $2827.4 \mu\text{m}^2$ |
| $g_{Na}$ | $50 \sim 60 \text{mS} \cdot \text{cm}^{-2}$ |
| $g_K$ | $5 \sim 100 \text{mS} \cdot \text{cm}^{-2}$ |
| $g_L$ | $0.033 \text{mS} \cdot \text{cm}^{-2}$ |
| $V_{Na}$ | $60 \text{mV}$ |
| $V_K$ | $-90 \text{mV}$ |
| $V_L$ | $-70 \text{mV}$ |
| $i_0$ | $19.1 \mu\text{A} \cdot \text{cm}^{-2}$ |
| $t_0$ | $1 \text{s}$ |

**Table 2.**
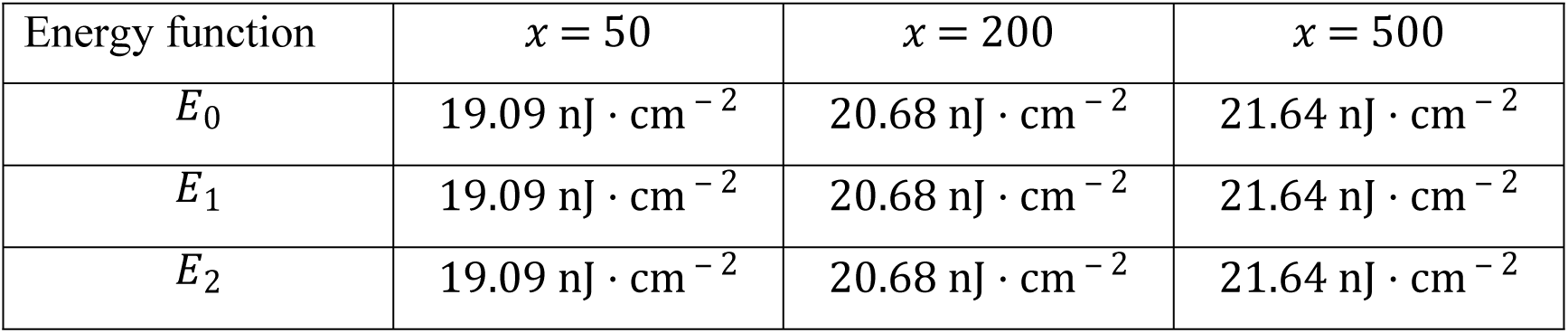
Energy dissipated by AP at three different sites.

| Energy function | $x = 50$ | $x = 200$ | $x = 500$ |
| --- | --- | --- | --- |
| $E_0$ | $19.09 \text{ nJ} \cdot \text{cm}^{-2}$ | $20.68 \text{ nJ} \cdot \text{cm}^{-2}$ | $21.64 \text{ nJ} \cdot \text{cm}^{-2}$ |
| $E_1$ | $19.09 \text{ nJ} \cdot \text{cm}^{-2}$ | $20.68 \text{ nJ} \cdot \text{cm}^{-2}$ | $21.64 \text{ nJ} \cdot \text{cm}^{-2}$ |
| $E_2$ | $19.09 \text{ nJ} \cdot \text{cm}^{-2}$ | $20.68 \text{ nJ} \cdot \text{cm}^{-2}$ | $21.64 \text{ nJ} \cdot \text{cm}^{-2}$ |

For the sake of the readers, we provide the deduction from Eq. [8] to Eq. [9]. From Eq. [8], we have
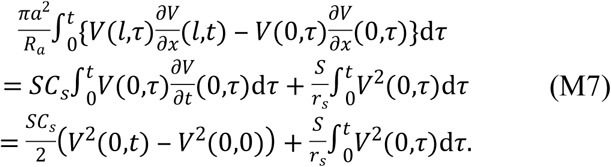

## Acknowledgments

ZJ was supported by the Fundamental Research Funds for the Central Universities under grant No. 3102017OQD073. YY thanks for the support from the National Natural Science Foundation of China (81761128011, 31571070), Shanghai Science and Technology Committee support (16410722600), the program for the Professor of Special Appointment (Eastern Scholar SHH1140004) at Shanghai Institutions of Higher Learning, and Omics-based precision medicine of epilepsy entrusted by the Key Research Project of the Ministry of Science and Technology of China (Grant No. 2016YFC0904400) for their support.

## ADDITIONAL INFORMATION

The authors declare no competing financial interests.

